## Supplemental Figures and Notes for "Contrasting Effects of Western vs. Mediterranean Diets on Monocyte Inflammatory Gene Expression and Social Behavior in a Primate Model"

### Supplementary Materials

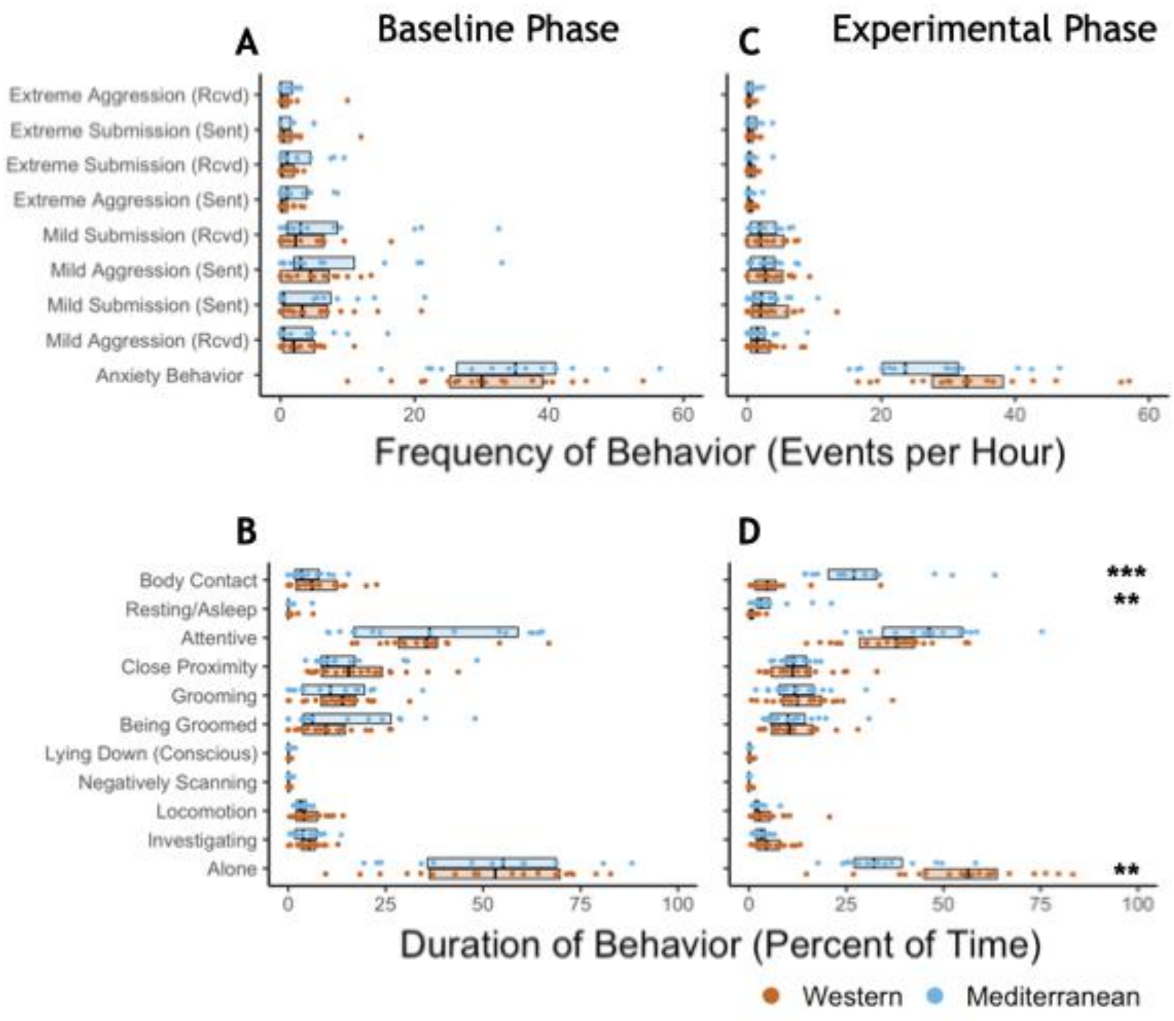


**Supplementary Figure 1. Diet manipulation altered behavior. (A) Rates and (B) duration of behaviors during the baseline phase, prior to diet manipulation.** There were no differences in behavior between the Western- and Mediterranean-fed groups. **(C) Rates and (D) duration of behaviors during the experimental phase.** The boxplots depict the per-group medians and interquartile ranges for each behavior. Animals fed the Western diet are colored orange, and those fed the Mediterranean diet colored blue. Significant differences in observed behavior between the diet groups during the experimental phase are indicated (Mann-Whitney U test, Holm-Bonferroni adjusted $p < 0.05$ *, $p < 0.01$ **, $p < 0.001$ ***)

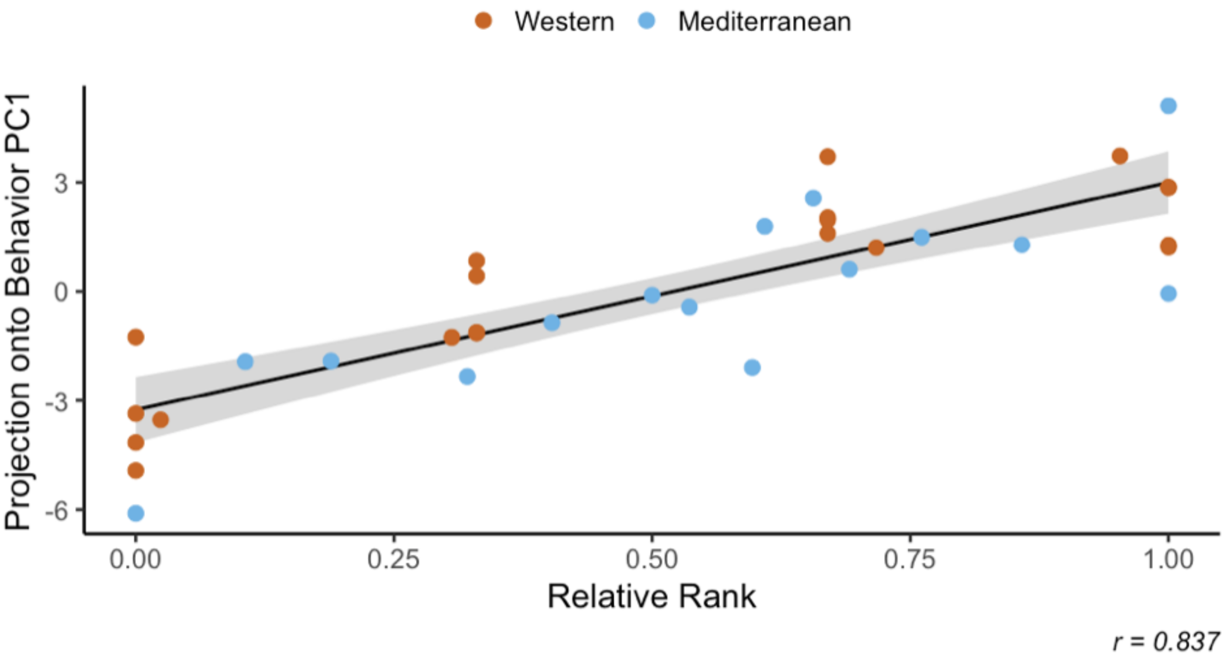


**Supplementary Figure 2. The first PC of all behavioral data captures dominance rank.** The first axis of variance in behavior—which explained 31.2% of the overall variance—was significantly positively correlated with dominance rank across diets (Pearson's $r = 0.837$, $p = 3.9 \times 10^{-10}$). All monkeys are assigned a rank between 0 and 1 based on the outcomes of dyadic interactions, where a higher rank indicates more dominant social status.

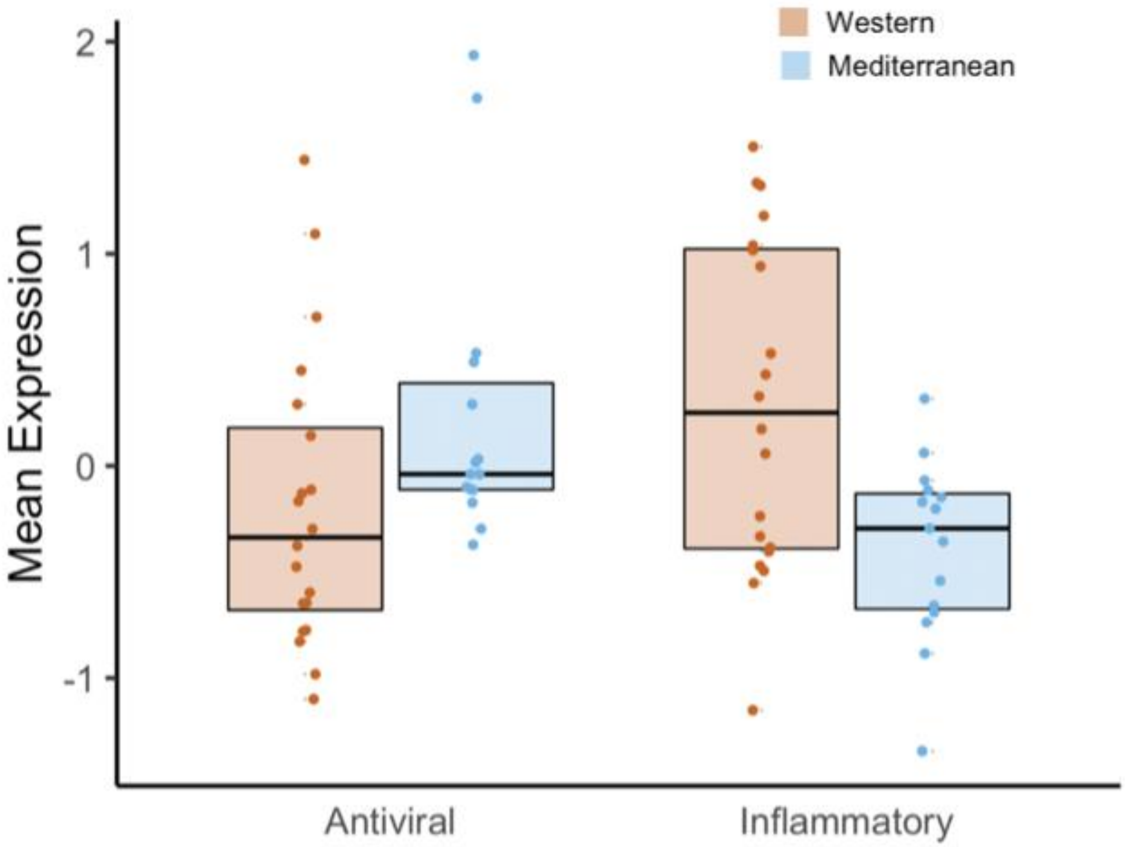


**Supplementary Figure 3. Expression of genes in the conserved transcriptional response to adversity (CTRA**(Cole et al., 2015)**) indicate inflammatory effects of a Western diet that parallel the effects of social adversity.** Western-diet fed animals exhibited significantly higher expression of pro-inflammatory genes involved in the CTRA (Mann-Whitney $U = 222$, $p = 0.016$), and lower expression of antiviral- and antibody-related CTRA genes (Mann-Whitney $U = 82$, $p = 0.023$). See Table S1 for CTRA categories.

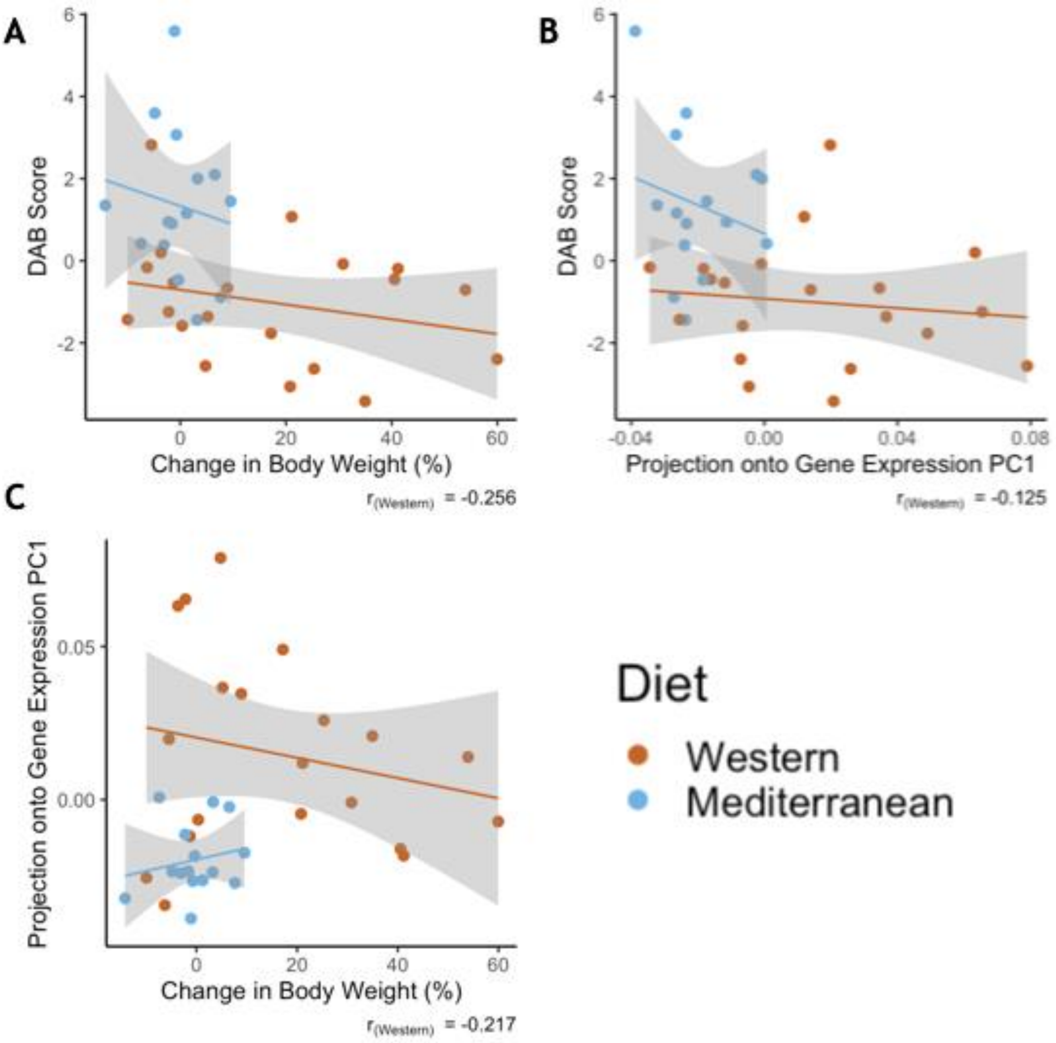


**Supplementary Figure 4. Greater phenotypic variability in Western diet fed monkeys does not show consistency in individual responsiveness across phenotypes. A)** Monkeys fed the Western diet showed more variability than monkeys fed the Mediterranean diet in both diet-altered behavior (DAB) and change in body weight. However, the two phenotypes were not correlated within monkeys fed the Western diet (Pearson's $r_{Western}$ = -0.256, $p$ = 0.28). **B)** Western fed monkeys also showed more variability in the first principal component (PC1) of gene expression than Mediterranean fed monkeys. DAB and PC1 of gene expression were not significantly correlated in Western fed monkeys (Pearson's $r_{Western}$ = -0.125, $p$ = 0.60). **C)** PC1 of gene expression and change in body weight were not significantly correlated in Western fed monkeys (Pearson's $r_{Western}$ = -0.217, $p$ = 0.36).

9

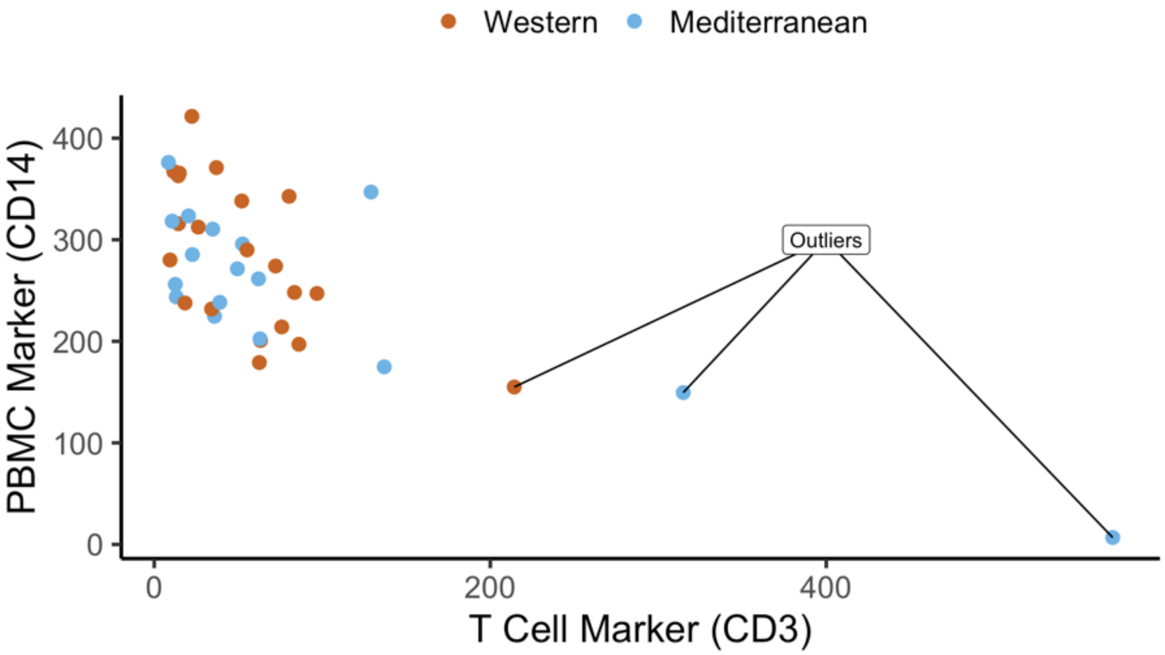


**Supplementary Figure 5. Quality control of cell purity by CD14 and CD3 expression levels: three samples were excluded due to lower CD14 and high CD3 – possible T cell contamination.** Normalized expression (reads per kilobase million) of CD14 and CD3 are plotted as markers of monocytes and T cells, respectively. Three samples were excluded as outliers due to possible T cell contamination.

10

11

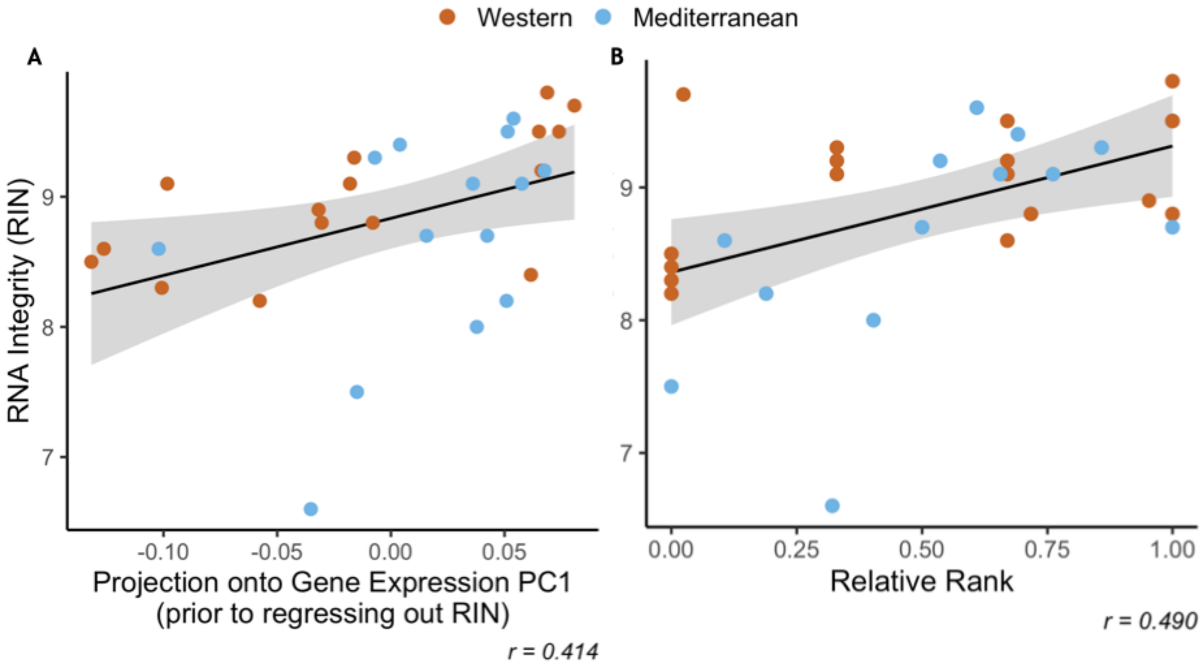


**Supplementary Figure 6. RNA Integrity was correlated with both uncorrected gene expression and relative rank. A)** RNA integrity (RIN) was correlated with PC1 of gene expression (62.4% of overall variance in gene expression) prior to correction for batch effects (Pearson's $r = 0.414$, $p = 0.020$). Because of this, RIN was included as a batch effect prior to downstream analysis. **B)** RIN was also correlated with relative dominance rank (Pearson's $r = 0.490$, $p = 5.1 \times 10^{-3}$). Points are colored to indicate Western (orange) or Mediterranean (blue) diet to show that diet did not have an interactive effect in either case.

12

**Supplementary Note 1. Regarding rank and RNA integrity (RIN).**

Because of the well-established effects of social status on atherosclerosis(Addo et al., 2012; Hallman et al., 2001; Rosengren et al., 2004; Shively et al., 1990; Steptoe & Kivimäki, 2012, 2013; Stuller et al., 2012; Yusuf et al., 2004), inflammation(Kiecolt-Glaser, 2010; Maes et al., 1998; Steptoe et al., 2007), and immune cell gene regulation(Brydon et al., 2005; Chen et al., 2008, 2008, 2011; Cole, 2013; Miller et al., 2008; Snyder-Mackler et al., 2016; J. Tung et al., 2012; Jenny Tung & Gilad, 2013), one of the goals of this study was to examine of social status interacted with diet to alter monocyte gene regulation. Specifically, we hypothesized that the promotion of regulatory polarization from the Mediterranean diet would attenuate the deleterious effects of social subordination. As with many RNA-sequencing data, the RNA integrity (RIN) in our data set was correlated with the first axis of variance in a PCA analysis of gene expression (Pearson's $r = 0.414$, $p = 0.020$; Fig. S6A) and thus needed to be controlled for. Unfortunately, in this dataset, RIN was significantly correlated with dominance (Pearson's $r = 0.490$, $p = 5.1 \times 10^{-3}$; Fig. S6B). Thus, when we controlled for RIN prior to downstream analysis, we also removed our ability to detect the effect of dominance rank on gene expression. Our subsequent analyses, and sampling at later timepoints in the study, led us to conclude this correlation was purely due to random and unidentified technical reasons (and that dominance rank does not influence RNA integrity). Samples collected at other timepoints in this study will allow us to address the potential interactions between diet and social status.
